# Microxanox: an R package for simulating an aquatic *MICR*obial ecosystem that can occupy *OX*ic or *ANOX*ic states

**DOI:** 10.1101/2023.02.06.527266

**Authors:** Rainer M Krug, Owen L. Petchey

## Abstract

*Microxanox* is an R package to simulate a three functional group ecosystem (cyanobacteria, phototrophic sulfur bacteria, and sulfate-reducing bacteria) with four chemical substrates (phosphorus, oxygen, reduced sulfur, and oxidized sulfur) using a set of ordinary differential equations. Simulations can be run individually or over a parameter range, to find stable states. The model can be implemented with different numbers of species per functional group. The package is constructed in such a way that the results contain the input parameter used, so that a saved results can be loaded again and the simulation be repeated. Furthermore, the package framework and code should serve as a useful starting point for making simulation models of other types of ecosystem.

## 1. Required Metadata

### 1.1. Current code version

Ancillary data table required for subversion of the codebase.

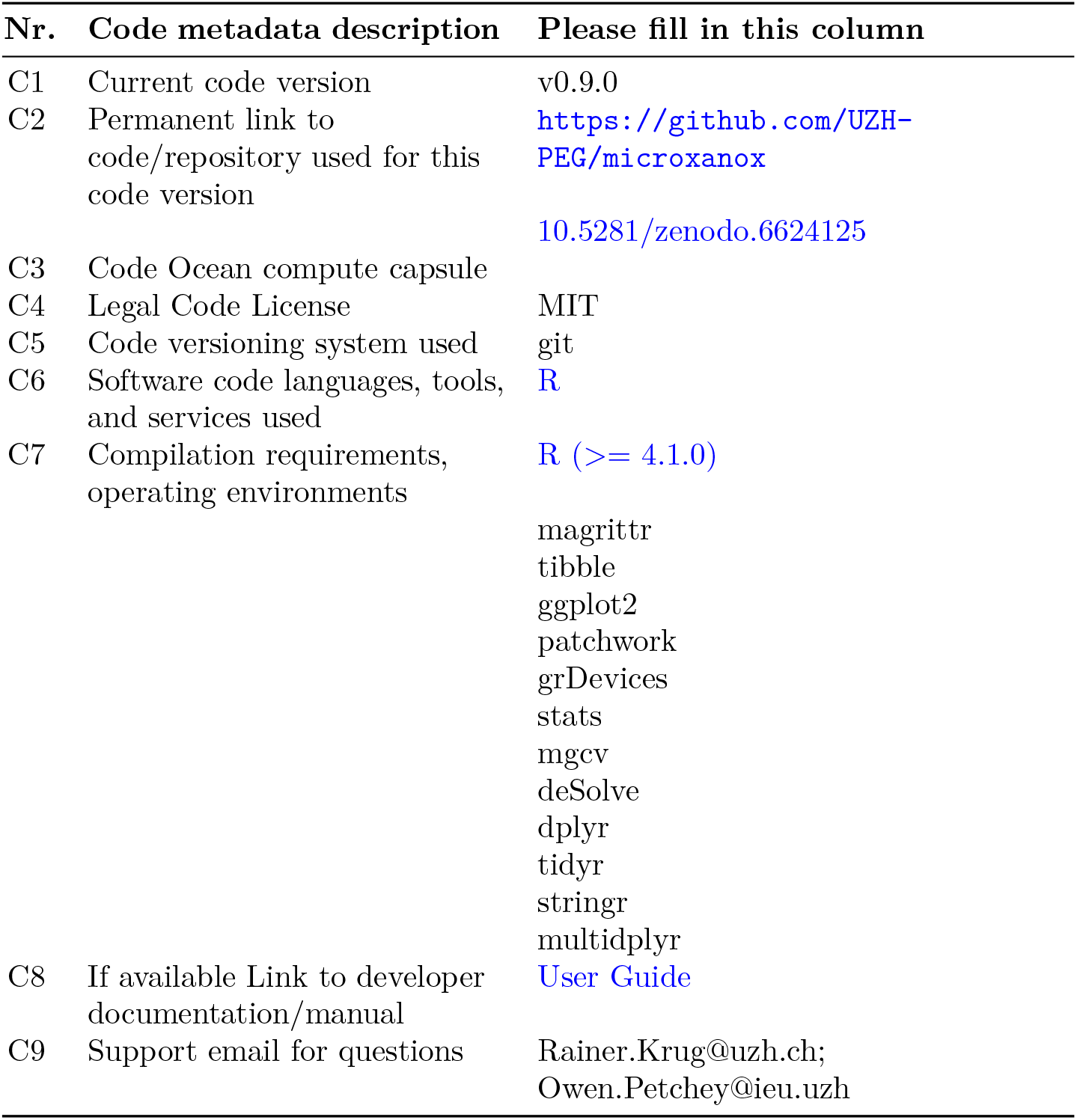

## 2. Motivation and significance

Mathematical models play a key role in the development of understanding about how ecosystems work and how they respond to environmental changes [1, 3, 14]. They are also important for developing hypotheses to test in empirical studies. One area of ecology in which models have played a influential role is how ecosystems respond to gradual change in an environmental driver [10]. An environmental driver is a environmental condition that affects an ecosystem, but is assumed to not be affected by the ecosystem, such as the rate of nutrient input into a lake.

It is conceivable that an ecosystem state, such as the total biomass of a particular type of bacteria, may remain unchanged when an environmental driver changes. It is also possible that the ecosystem state changes gradually. It is also possible that the ecosystem state changes abruptly to a new state that is difficult to recover from [10]. This possibility for abrupt, perhaps catastrophic changes that are difficult to reverse causes considerable concern [4, 7, 13].

An example where a gradual change of an environmental variable causes an abrupt change of the system is the switch from an aerobic (oxygen is available for metabolism) to anaerobic (oxygen generally unavailable) state in a microbial ecosystem. This system has been investigated by Bush et al. [2] in a simulation study of a mathematical model. Three types of microbes occur in the model: cyanobacteria (CB) dominating the oxic state, and two types of sulfur bacteria that dominate the anoxic state (sulfate reducing bacteria (SB) and phototrophic sulfur bacteria (PB)). The model shows that gradual change in the rate at which oxygen could diffuse into the ecosystem (termed the oxygen diffusitivity) could cause catastrophic changes in the ecosystem state that would be difficult to reverse.

One feature of the study by Bush et al. [2] was limited biodiversity. Specifically, there was no biodiversity within each of the three types of bacteria. This leaves open the question of if and how biodiversity within these types (i.e. functional groups) of bacteria affects the ecosystem response to environmental change. This limitation is not specific to the study of Bush et al. [2]. There are few if any studies of the effects of biodiversity on abrupt transitions between ecosystem states.

We decided to fill this research gap by making a simulation study of how within functional group biodiversity affects ecosystem responses to environmental change Limberger et al. [6], and to base our work on the work and model of Bush et al. [2]. It was with this goal in mind that we developed the *microxanox* package [5]. The first stage of development was to write code from scratch (as there was no available code to start from) and to confirm that this new implementation would reproduce the previously published results. The resulting reproduction is available as one of the package vignettes: [vignette Partial reproduction of Bush et al](https://uzh-peg.r-universe.dev/articles/microxanox/partial-reproduction-Bushetal2017.html).

The second stage was to add functionality that would be necessary to answer our research question. Most importantly, we made it possible to have multiple species of bacteria within each of the three functional groups, for the multiple species to differ in their characteristics, and to vary the number of species and amount of variability among them. We also added functionality that allowed: temporally varying environmental conditions, addition of random noise to state variables, and immigration. In addition to the model itself, the package includes some functions to analyse the results as well as to visualize the results to provide a starting point for customized visualizations based on own requirements. The basic and additional functionality is described in the package [User Guide](https://uzh-peg.r-universe.dev/articles/microxanox/User-guide.html).

## 3. Software description

The *microxanox* package is for simulating a three functional group system (*CB*: cyanobacteria, *PB*: phototrophic sulfur bacteria, and *SB*: sulfate-reducing bacteria) with four chemical substrates (*P*: phosphorus, *O*: oxygen, *SR*: reduced sulfur, *SO*: oxidized sulfur). It includes feedback between organisms and biogeo-chemical processes and is based on Bush et al. [2] (See Bush et al. [2] for a detailed discussion of the model). At the core of the simulations is a set of ordinary differential equations (specified in the function `bushplus_dynamic_model()`, though this function need not be directly called). There are functions for running individual simulations and for running a set of simulations across, for example, a range of environmental conditions.

To make the simulation run with multiple species per functional group, we expressed different species characteristics in the elements of vectors and matrices. We also coded the ordinary differential equations to include the vectors and matrices, and to use matrix mathematics. In this way, we made it possible to run simulations with different numbers of species without having to change the underlying code.

The package functions and code have modular structure so that new functionality can be easily added. E.g. temporally defined events of any type could be specified. Further, all parameter values required to run a simulation are stored in one object. Lastly, the general structure of the code should make it straightforward to adapt the model to other similar systems (described in more detail in the Impact section).

### 3.1. Software architecture

The framework used when writing this package aimed to maximise simplicity for the user, and to make it straightforward to reproduce results (see the supplement [8] to Limberger et al. [6] for an example of how this is used). As such, all the parameters needed to run a simulation or find a stable state (i.e. the final state of the ecosystem) are contained in a single object (which can easily be created using included functions). This parameter object is given to a function that runs the simulations and returns the results. The returned results object is identical to the parameter object but with an additional slot named `results`, which contains the results of the run. Thus the returned results object contains the simulation conditions (parameters) as well as the results, and can be used to run the simulation again. This promotes reproducibility and makes incremental changes of individual parameters and re-running the simulations straightforward.

In the following sections we describe how to use the package to run one simulation and to find steady states across an environmental gradient.

### 3.2. Running one simulation

A typical simulation would look as shown in Figure 1.

**Figure 1:**
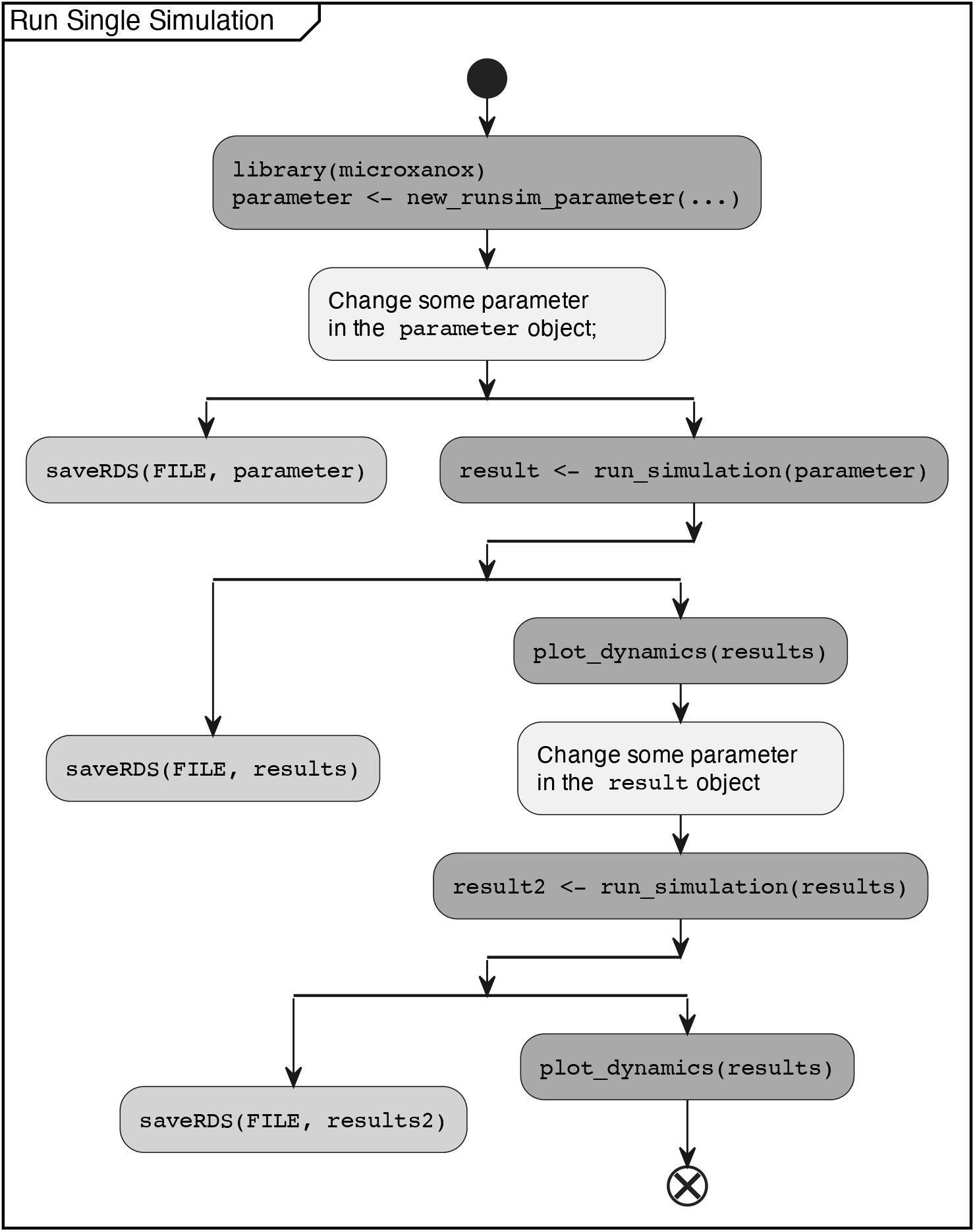
Typical flow of a simulation. Dark Grey boxes: commands necessary for simulation; Light Grey:Saving of parameter and results; Lightest Grey: Different non specified commands.

A simulation is run using the `run_simulation()` function. In this function, the ODEs are solved using the function `ode()` in the package *deSolve* package [12]. The `run_simulation()` function needs only one argument - an object as created by the function `new_runsim_parameter()`. The parameter object returned by `new_runsim_parameter()` contains among other things the strain_parameter object, which can be created by the function `new_strain_parameter()`. For a detailed description of the parameter objects, their meaning and how they are created and have values set and changed please see the *User Guide* which accompanies the package or is available at [User Guide](https://uzh-peg.r-universe.dev/articles/microxanox/User-guide.html).

After the parameter object has been defined, it can be used in the `run_simulation()` function. The function returns an object which is identical to the parameter object, except of an additional slot containing the results. This design produces a fully reproducible object as it can be used as a parameter object to be fed back into the `run_simulation()` function to run the simulation again from the parameter used to generate the results before.

### 3.3. Finding Stable States

The general approach used to find the stable state of the system with a specific parameter set is to run the simulation for a long time and record the final state. When one does this across a range of environmental conditions, one discovers how the steady state of the system responds to the environmental conditions. The package contains functionality for finding steady states that correspond to values of one environmental driver, namely the value of oxygen diffusivity.

Two methods for finding steady states are implemented. The first runs a separate simulation for each combination of starting conditions and oxygen diffusivity (we term this the *Replication method*). This is the method used in the Bush et al. [2] study. The second runs two simulations, one with step-wise and slowly temporally *increasing* oxygen diffusivity, and the other with step-wise and slowly *decreasing* oxygen diffusivity. During this temporal environmental change, the state of the system is recorded just before change to a new oxygen diffusivity (we term this the *Temporal method*). We implemented two methods since there is no definitive best method, and in order to check if results were sensitive to choice of method.

The replication method is implemented in the function `run_replication_ssfind()` which takes a parameter object as returned by the function `new_replication_ssfind_parameter()` and the number of cores for multithreading the simulation. As the multithreading uses the package function `mclapply()` from the package `parallel` [9], the multithreading only works on Linux and Mac. It is planned to move to `parLapply()` [9] in a future release.

The temporal method is implemented in the function `run_temporal_ssfind()`, which takes a parameter object as created by the function `new_temporal_ssfind_parameter()`. It is planned for a later release, to run these two simulations in parallel.

For a more detailed walk-through of these two approaches and explanation please see the [User Guide](https://uzh-peg.r-universe.dev/articles/microxanox/User-guide.html).

### 3.4. Analysing and visualising results

From the results returned, summary measures about how the ecosystem stable states respond to environmental change can be extracted. The function get_stability_measures() returns quantities such as the amount of environmental change required for the system to abruptly change to a different state.

The function plot_dynamics() plots a single simulation run, as returned from the run_simulation() function. This function is only provided as a convenience function to provide a way to easily see the results of a simulation run. An example plot resulting from this function is shown in Figure 2.

**Figure 2:**
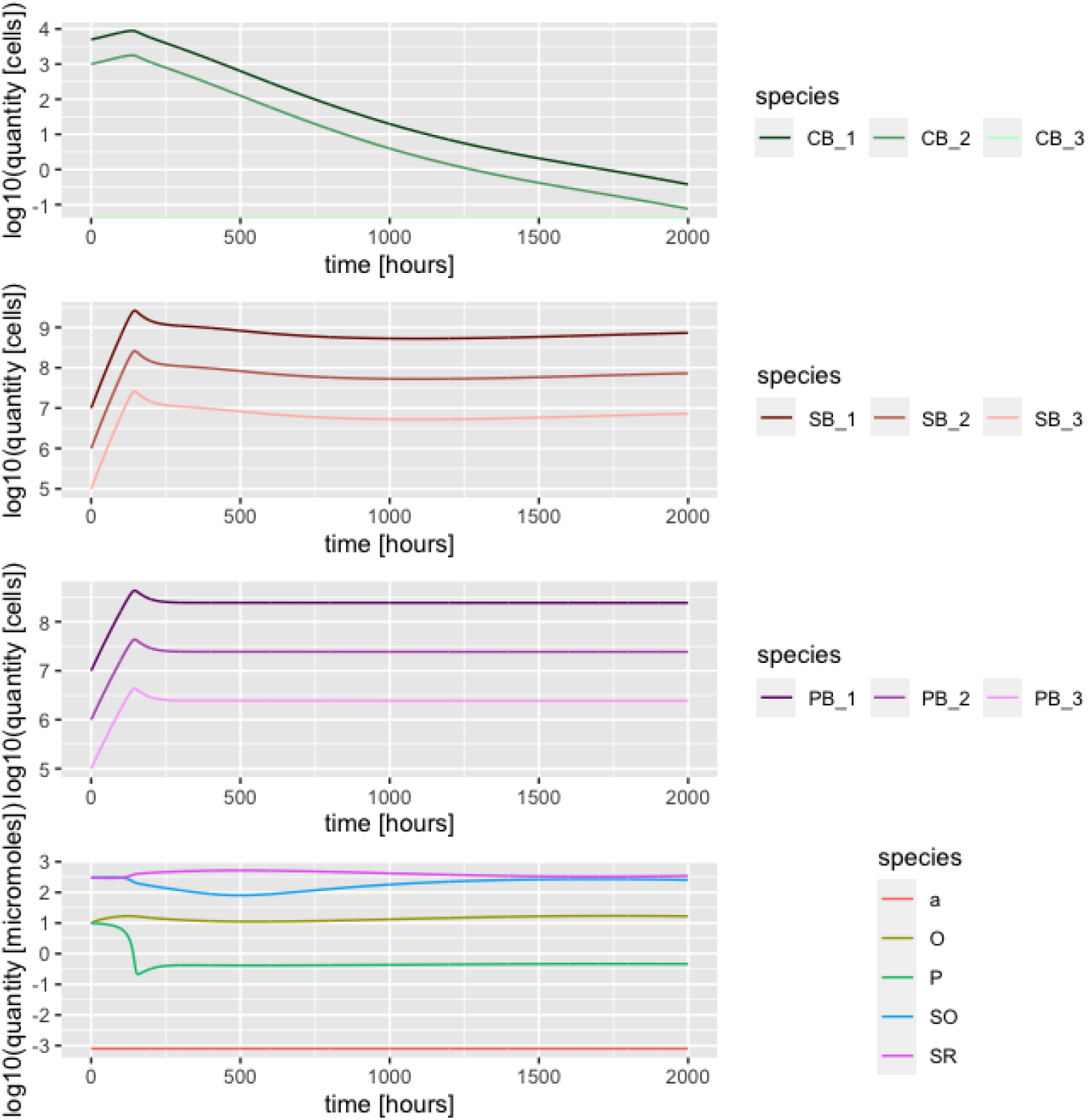
Results of a simulation run shown using the function *plot*_*dynamics*(). In this case, there were three strains per functional group, though strains within functional groups had identical properties in this example. CB_1 = cyanobacteria strain 1; SB_1 = sulfur reducing bacteria stain 1; PB_1 = phototrophic sulfur bacteria strain 1.

## 4. Impact

The open source implementation and extension of the model used in Bush et al. [2] provides the means of reproducing the results published while at the same time provides the means of doing unique, innovative, and important investigations of how ecosystems respond to environmental change, and how biodiversity may modulate this response.

The design of the package code and functionality is with reproducibility in mind: the combination of all parameters being in a single parameter object as well as the return of the simulation as a result object which inherits from the parameter object provides a relatively easy to use framework to implement reproducible experiments.

Here we evidence the impact of the *microxanox* package by describing three use cases and then by describing how the package can be a starting point for models of other ecosystems. The first two use cases are described in detail (including the code for reproducing them) in the *User Guide* and the *Partial Reproduction* vignettes. The third is taken from Limberger et al. [6] and Petchey et al. [8].

### 4.1. Use case 1: Regime shifts during temporal environmental change

The study of Bush et al. [2] includes simulations of the effect of oxygen diffusivity (an environmental driver, in the sense that it affects the ecosystem but is not affected by it) on the ecosystem state (oxic or anoxic). The *microxonax* package contains functionality to make a specific temporal pattern of change in the oxygen diffusivity. As well as allowing individual simulations during which oxygen diffusivity varies, this functionality forms the basis of the temporal method for finding stable states.

An example of this functionality is given in the *Partial Reproduction* vignette, which we briefly show here (Figure 3). The example is composed of a single simulation, at the beginning of which the system is in the oxic state with high abundance of cyanobacteria. Oyxgen diffusivity is then slowly decreased and eventually, around hour 30’000 the system switches to the anoxic state, with high abundance of both sulfur bacteria types. The oxygen diffusivity is then increased and at around hour 38’000 the system abruptly switches back to the oxic state.

**Figure 3:**
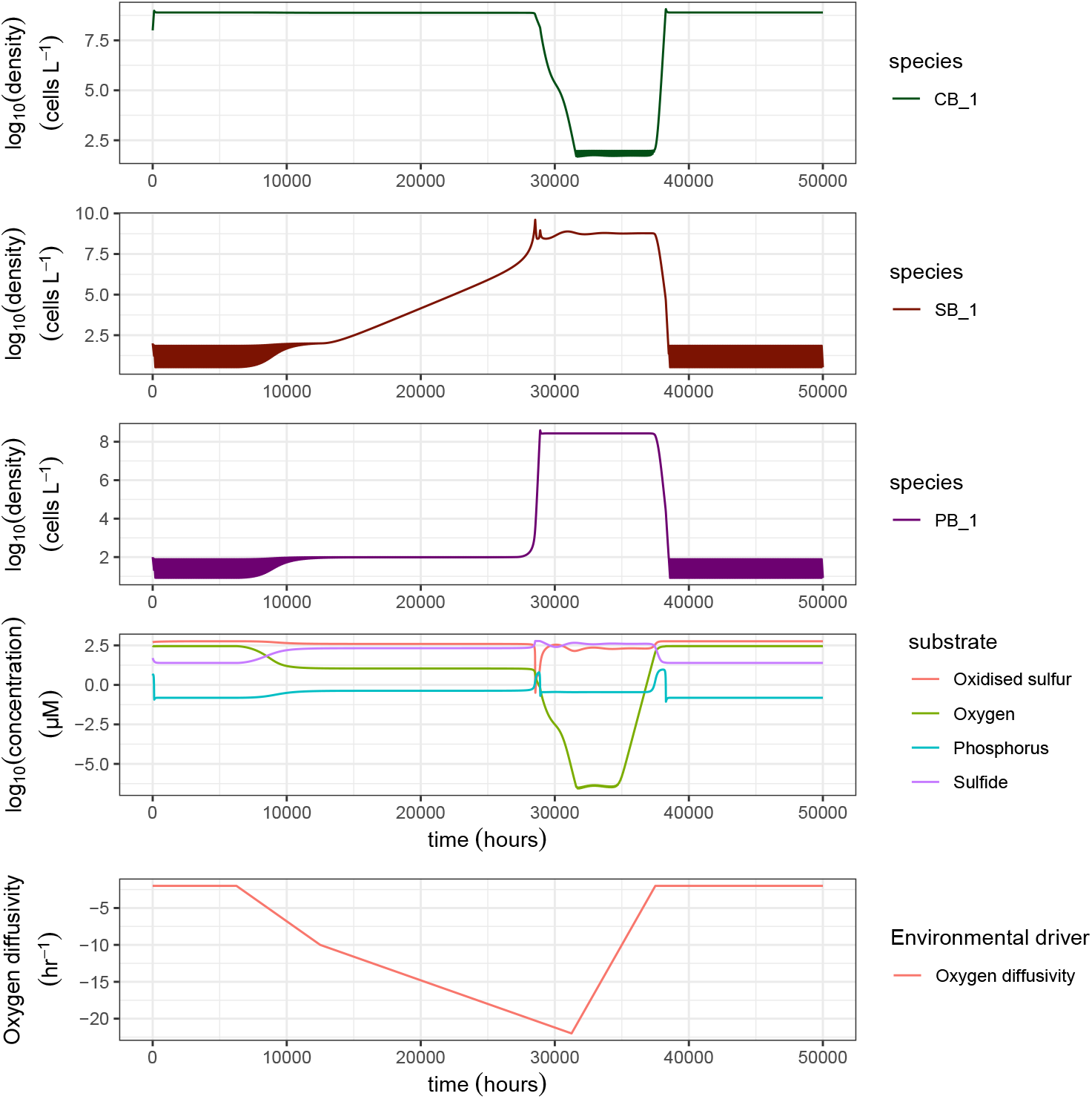
The temporal dynamics of the ecosystem model when an environmental condition (here parameter *a*, the oxygen diffusivity) changes. Plot of the stable states of the simulation runs under different oxygen diffusivity. In this simulation there is only one strain in each functional group. CB extunderscore1 = cyanobacteria strain 1; SB_1 = sulfur reducing bacteria stain 1; PB = phototrophic sulfur bacteria strain 1. Here we show a figure adapted from the output of the *plot*_*dynamics*() function.

Also visible in the results are thick lines showing abundances of bacteria when abundances are low. This is due to the implementation of a function that at regular intervals, increases the abundance to a preset level. This prevents abundances reducing to very small numbers. The function that implements this increase abundance can also be made to add a certain abundance to each strain at regular intervals, thus simulating immigration in to the system.

### 4.2. Use case 2: The extent of hysteresis depends on community composition

The package contains a function to extract summary features of ecosystem responses to environmental change, such as the amount of hysteresis displayed by the ecosystem. Hysteresis is a key feature of ecosystem responses to environmental change, because it is related to how difficult it can be to reserve the effects of environmental change [10]. The amount of hysteresis is measured as the extent of the environmental condition (here oxygen diffusitivity) for which there were two stable states. I.e. it is the extent of the environmental conditions for which historical conditions play an important role in determining the current system state (a definition of hysteresis).

Using the package to calculate the extent of hysteresis involves setting ecosystem and simulation parameters, including parameters for the finding of stable states across an environmental gradient, running the stable state finding function, and analysing the results with the function that calculates extent of hysteresis. The code for this is provided in the *User Guide*.

The results show that the amount of hysteresis depends greatly on the combinations of organisms present (Figure 4). For example, with only the CB (cyanobacteria) present, there was no hysteresis. In contrast, the presence of both CB and SB (sulfate reducing bacteria) led to a large amount of hysteresis. (These results are also given in the *Partial Reproduction* vignette.)

**Figure 4:**
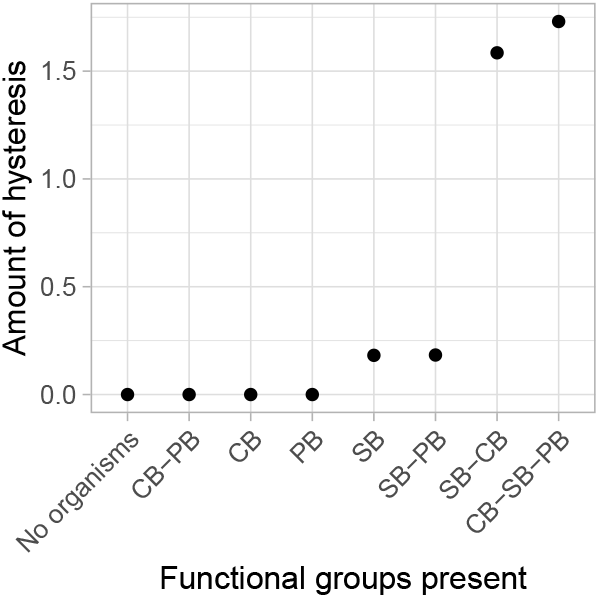
The amount of hysteresis depends on the combination of types of organisms present. The model is entirely deterministic, hence there are no error bars.

### 4.3. Use case 3: Effects of functional diversity on regime shifts

As discussed in the Introduction section, the package was motivated by the question of how biodiversity influences ecosystem responses to environmental change. Extensive results concerning this question are given in a separate publication Limberger et al. [6]. Here we describe one of the results, which is that having biodiversity in a functional group can allow state changes to occur that otherwise would not have. I.e. biodiversity can qualitatively change the state of the ecosystem.

Biodiversity was added to the functional groups using the new_strain_parameter() function to create a parameter set with multiple species per functional group (albeit all with identical features) and then to add variability among the species by calling the add_strain_var() function. This function takes an already existing parameter set and adds the specified about of variation. The new parameter object is then used as before.

Figure 5 shows a simulation with two species (strains) in each of the three functional groups. The ecosystem starts in the oxic state, though with relatively high abundance of each functional group. The strain of SB that is more tolerant to oxygen (SB_1) initially decreases in abundance, but then increases, and the other (SB_2) strain then becomes abundance and SB_1 declines. Furthermore, the cyanobacteria crash in abundance, and the system switches to the anoxic state. In contrast, if there are two identical strains with tolerance half way between those in Figure 5 the ecosystem remains in the oxic state.

**Figure 5:**
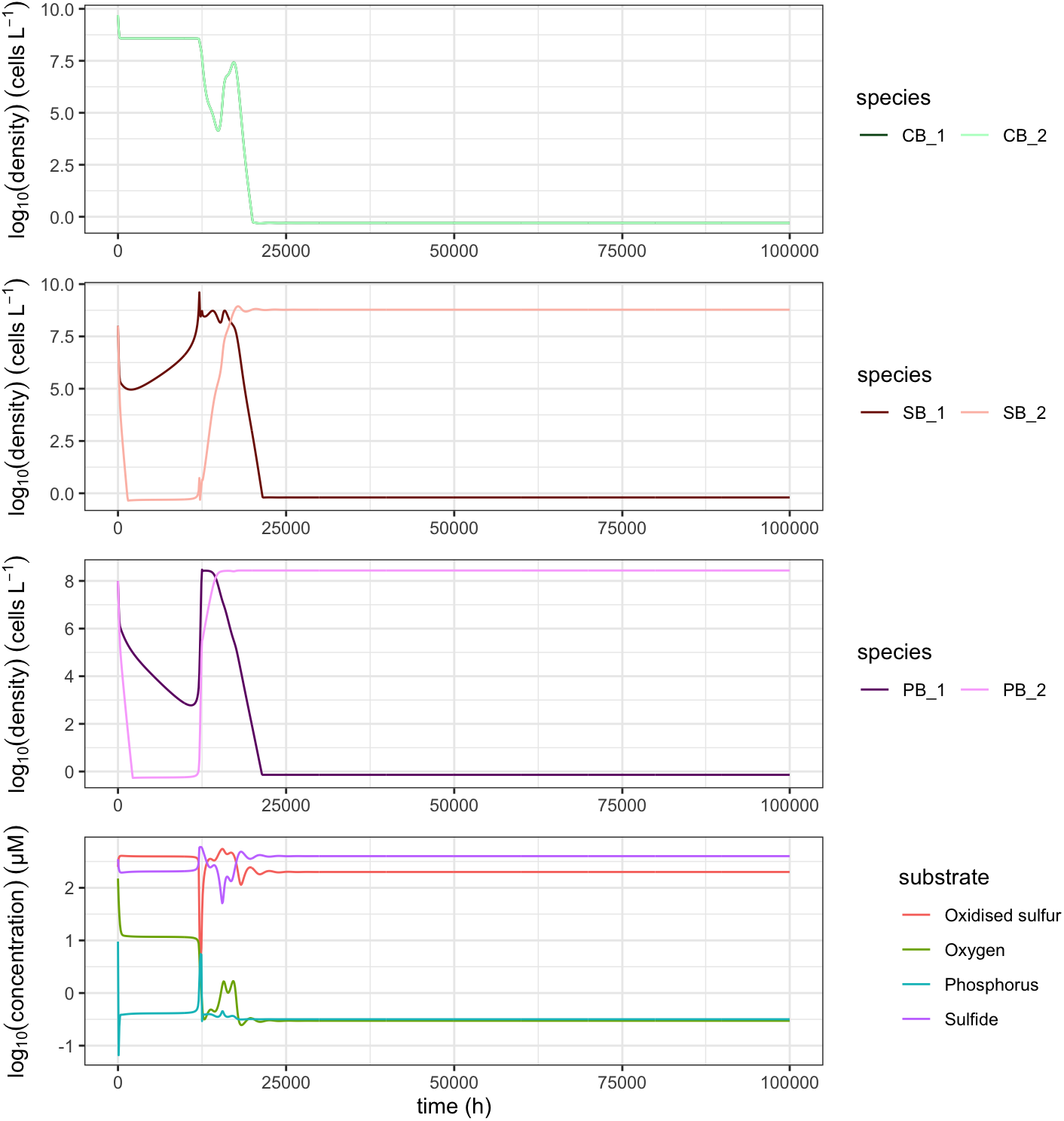
The dynamics of the ecosystem when there are two species in each functional group, and some variation (diversity) in species parameters.

### 4.4. Adapting for other ecosystems and organisms

We anticipate that the package can be a useful starting point for investigating models of other types of ecosystem and how biodiversity in them affects responses to environmental change. The overall framework of the package, the purpose of each function, and the objects used for storing parameters and results could be retained. E.g., all such models would have parameters that differ among species and need to be described in an object, and studies will often need to run simulations and sets of simulations across environmental conditions.

Researchers wanting to model a new ecosystem do not, therefore, have to start from scratch. This will relieve researchers from needing to make software design decisions, and rather focus on appropriately representing their ecosystem and finding the results that interest them. Nevertheless, adaptation of the code in the package will require a person / persons that can take a conceptual model of an ecosystem and then represent that in terms of parameters and rate equations, and that is relatively proficient in R programming.

## 5. Conclusions

The *microxanox* R package allows the simulation, visualisation, and analysis of a model of a microbial ecosystem while allowing variation in the amount of diversity containing in each of the functional groups of organisms present. It has been used for the research described in another paper that describes one of the first investigations of the effects of diversity on ecosystem resilience Limberger et al. [6]. In that paper, we show that diversity can have large and important effects of ecosystem responses, highlighting the need for models such as ours, with which one can easily manipulate the amount of biodiversity. The *microxanox* package has also been used to reproduce the results of the paper that inspired the package development [2].

The package greatly lowers the amount of work required in further investigations of the specific ecosystem modelled. There has, for example, been quite limited investigation of how biodiversity influences the short-term responses of the modelled ecosystem to environmental change. Likewise, the package could be used to power an investigation of the effects of biodiversity on the usefulness of early warning signals of abrupt ecosystem change [11]. In addition this package could be used as a template for the implementation for developing models of other types of ecosystems and organism. By doing so, other models can profit from the overall framework used, and the reproducibility aspects as well as the flexibility implemented.

## 7. Acknowledgements

This project was part of SNF Project 310030_188431. The project was also supported by the University of Zurich Research Priority Programme in Global Change and Biodiversity.

